# Calling site preference reduces masking interference of acoustic signals among sympatric bush frogs and facilitates coexistence

**DOI:** 10.1101/2023.09.25.559099

**Authors:** A V Abhijith, Shomen Mukherjee

**Affiliations:** Jawaharlal Nehru Centre for Advanced Scientific Research (JNCASR), Rachenahalli Lake Rd, Jakkur, Bengaluru, Karnataka 560064, India; Azim Premji University, Survey No 66, Burugunte Village, Bikkanahalli, Main Road, Sarjapura, Bengaluru 562125, Karnataka, India; Biological and Life Sciences Division, School of Arts and Science, Ahmedabad University, Ahmedabad, Gujarat 380009, India; Ecology, Environment and Climate Change Research Cluster, Ahmedabad University, Ahmedabad, Gujarat 380009, India

**Keywords:** Acoustic Niche Hypothesis, species coexistence, Rachophoridae, cocktail party problem, coffee plantation, Western Ghats

## Abstract

The degree of signal overlap among sympatric species strongly influences the efficiency of intraspecific communication. This phenomenon is particularly pronounced among species that rely heavily on acoustic communication for locating mates and defending territories. Signal interference presents the classic cocktail party problem and, hence, can pose challenges for these species as it can likely hinder the detection, recognition, and localisation of conspecifics. To overcome this challenge, sympatric species are expected to partition their acoustic space to minimise masking interference. We studied a group of sympatric and closely related endemic frogs in coffee plantations of the Western Ghats, India. This community is active for a limited period during the monsoon and, therefore, has a restricted breeding season. Acoustic communication during the breeding season is integral to these frog species, making them an ideal system to investigate possible strategies adopted by a community to minimise masking interference. Using observational data on the vertical height of calling sites and acoustics data, we show that the frog assemblage can partition in multidimensional trait axes, such as call frequency and space. In addition, a novel coexistence strategy emerges when combining these results, where frogs with similar acoustic parameters partition more in their space use. Our results broadly suggest that competition for acoustic space can drive signal and space-use partitioning and that vertical call site selection can enhance the minimisation of masking interference. Our study is among the few studies that test *spatial stratification* in conjunction with *spectral stratification* among coexisting species, and to the best of our knowledge, the first on amphibians.

## INTRODUCTION

Auditory or acoustic signals are an important mode of communication in a wide range of organisms, e.g., mammals, amphibians, birds, and insects. They are important for locating mates (Bailey & Zuk 2008), in sexual selection decisions (McGregor 2005), and finding competitors (McGregor & Peake 2000), hence affecting an individual’s fitness. Social aggregation of multiple vocalising species (i.e., a noisy environment) can challenge the sender and receiver of such signals by disrupting their quality via masking interference, which is analogous to the human cocktail party problem (Bee & Micheyl 2008). So, how do organisms address such a challenge?

Studies of the cocktail party problems have received less attention despite it being common in nature (Hulse 2002). Information about how animals address this problem is vital for understanding the evolutionary diversity in sensory mechanisms, receiver psychology, applications such as network theory, and the effects of anthropogenic noise on wildlife (Bee & Micheyl 2008). This study used an arboreal frog community to understand how amphibians address the cocktail party problem.

Among many features that characterise frogs, their distinct vocalisation and acoustic communication constitute an important and conspicuous part of their breeding biology. Males almost exclusively give out these calls during the breeding season to either attract mates, defend territory or compete with other neighbouring males (Wells 2007). The advertisement calls are crucial for male success as females base mate choice decisions on the acoustic properties of these calls (Sullivan *et al*. 1995). Apart from a few exceptions, most species call in aggregate, especially in the tropical regions known for their amphibian diversity (Duellman 1967b, Schwartz & Wells 1983).

Sympatric frog species calling simultaneously can lead to two problems for intraspecific communication (Wells 2007). First, the calls of other species may mask and obscure the features of the calls meant for conspecifics. Second, if calls of sympatric species are very similar, females might get confused and may approach or choose males of the wrong species. Given the above challenges of aggregate calling, closely related sympatric species are expected to have evolved certain strategies to coexist and maintain reproductive isolation, thus maximising their fitness.

Both competition theory (Lotka 1925, Volterra 1927) and empirical evidence (e.g., Stein *et al*. 2013) suggest that coexistence among species is possible when there is low interspecific competition. Ecologically, this means that coexistence is achieved mainly through resource partitioning (Colwell & Fuentes 1975, Diamond 1978). From an acoustics perspective, these strategies have primarily been approached from the Acoustic Niche Hypothesis (ANH) predictions (Krause 1987). ANH predicts minimal signal overlap among co-existing species to facilitate intraspecific communication. Despite many attempts at collating evidence supporting this hypothesis, the theory remains equivocal (Memet *et al*. 2022).

Since interactions between a signaller and receiver are typically not dyads but can involve multiple receivers (targeted or non-targeted), signalling studies have used the communication network perspective to understand the evolution of signals (McGregor & Peake 2000, McGregor 2005). If selection pressure from non-target individuals is high (e.g. from eavesdropping predators, see Peake 2005), unlike ANH’s, one predicts more signal overlap among sympatric species than expected by chance - the Network hypothesis (NH; Legett *et al*. 2019).

In the context of frogs calling in aggregation during mating season, intraspecific communication should be prioritised, especially to defend territories and to recognise potential male competitors and mates. This would be more pronounced in frog systems where breeding is seasonal (e.g., 2-3 months of monsoon season in India). Therefore, an investigation into the acoustic dynamics of closely related sympatric species during breeding season is expected to be synergistic with the predictions of ANH.

The ANH presents a gateway to comprehending the partitioning of the acoustic space based on the three dimensions of the acoustic niche (Littlejohn 1977). The first could be through *spatial separation*. This could be through the aggregation of individuals into single-species assemblages or by using different microhabitats for calling. Second, coexisting species may show *spectral stratification*, wherein each species uses a non-overlapping, unique frequency bandwidth for communication/signalling. Third, there could be *temporal partitioning* between species. This could either be through species-specific breeding seasons, or by each species having a unique time during its daily activity cycle to signal/call. Finally, coexisting species may have different coding patterns in their advertisement calls. This helps minimise mistakes in species identification but would have minimal effect on acoustic interference. Behavioural and community ecology studies have established that one or more of the abovementioned mechanisms may aid in reproductive isolation and thus allow the coexistence of closely related sympatric species (Duellman 1967a, Grafe 1996).

While studies have extensively tested the ANH in natural habitats such as forests, grasslands, ponds, etc. (e.g. Chek *et al*. 2003, Both & Grant 2012, Villanueva-Rivera 2014, Lahiri *et al*. 2021), few have tested how sympatric species had adapted to novel (human-made) habitats to avoid the cocktail part problem. Such studies are vital for conserving species in human-dominated landscapes. This study examined the spatial and spectral partitioning among four closely related sympatric species of male bush frogs coexisting in coffee plantations in the southern part of the Western Ghats Mountains (a global biodiversity hotspot) in south India. These frogs, which coexist in the evergreen forests of the southern Western Ghats, a biodiversity hotspot, have evolved mechanisms to partition a condensed micro-habitat (∼300cm high coffee plants) to avoid interference in their breeding signals.

The study is also novel because this is the first study which tests *spatial stratification* in conjunction with *spectral stratification* among coexisting species of amphibians. This is important because, unlike signal properties of species, which are constrained by morphology (Gillooly & Ophir 2010, Memet *et al*. 2022), most vocalising species are expected to be more flexible in selecting calling sites to enhance the efficiency of signal transmission (Wells 2007, Jain & Balakrishnan 2012, Chitnis *et al*. 2020). Hence, insights from this study are unlike most ANH studies (including ones in the wild), which primarily rely on only acoustics, therefore not testing for *spectral stratification* and *spatial partitioning* in the same system. Identifying and quantifying species-specific differences in space use for vocalising “daises” is essential to understanding the evolution of signal properties, both in the context of avoiding interference competition and in signal propagation.

## METHODS

The study was conducted in 5 coffee farms located within a 10km radius of Sulthan Bathery town, Wayanad, Kerala state, India, between May and July 2019. Post-selection, two 35m transect lines were laid in each farm. Only a single transect was laid in two farms because of size constraints. Three species of endemic bush frogs (*Pseudophilautus wynaadensis, Raorchestes akroparallagi, Raorchestes ponmudi*) are common in these farms, while a fourth species, *Raorchestes glandulosus*, is rare but present. Males of all four species of frogs use the coffee plants (all maintained within 300 cm height) for advertising (calling), finding mates, and copulation. Female frogs also seek mates on these plants, then lay eggs on the ground among leaf litter (Abhijith & Mukherjee 2020). All the four species are nocturnal.

### Assessing vertical stratification among sympatric frog species

On five farms, visual encounter surveys were conducted along the transects between 6–11 pm. When a calling frog was encountered, the species was noted, and the individual male’s exact height was recorded using a ruler. This data was then used to test for differences in calling heights among the four species.

### Assessing if observed vertical stratification patterns were a result of preference or interference

The observed vertical call site partitioning, if any, could be a result of an individual frog “preferring” a vocalising perch at a certain height or the individual is “restricted” to a height category because the other heights have already been occupied by conspecifics (or species). To determine which of these is true, surveys were conducted in five coffee farms between 6-11 pm between June and July 2019. Each farm was visited after rain when the frogs were acoustically in peak activity. Following a single call, an individual (male) was located on a plant, its species identification was confirmed, and its height measured. Following this, the rest of the plant was searched thoroughly (using a head torch) for other individuals, either of the same or other species. On average, 20 minutes were taken to search each plant thoroughly. This method was followed, and around ten per cent of the coffee plants on each farm were searched. This data was then used to understand patterns of species co-occurrence, i.e., the proportion of plants with single species, two and three species of frogs. A large proportion of plants with single species occurrence would signify “preference” for the height observed in the previously described vertical stratification study.

### Call frequency segregation among sympatric frog species

Five farms were visited after a good rain, a time when the frogs are acoustically in their peak activity. Five fifteen-minute audio recordings were made at random between 6–10 pm at each farm (one recording for each hour) using a ZOOM H2N multidirectional audio recorder. The recorder was mounted onto a Manfrotto tripod (approx. 5 feet from the ground) to avoid any disturbances during recording.

The audio recordings were then analysed using Raven Pro 1.6 software (Cornell Laboratory of Ornithology, Ithaca, NY, USA). This software provides a visual representation of the spectrum of frequencies of an audio signal as it varies with time, i.e., a spectrogram. The spectrogram was generated using a Hann window with a sample size of 510 samples and a 50% overlap between windows. Since each of the four species of bush frogs has a unique call, the spectrogram patterns for the calls of each of the four species were identified.

For each species, we labelled at least 65 notes using the cursor and selection box features available in Raven Pro. During the process, care was taken to digitise all note types to encompass the vocal repertoire of each species. In the case of species with multiple repertoires, we digitised all note types in comparable proportions to avoid over-representing a particular note.

We extracted the following eight parameters for all the digitised notes: Peak frequency (Hz), Peak Frequency Contour Max Frequency (Hz), Peak Frequency Contour Min Frequency (Hz), Average Entropy (bits), Centre Frequency (Hz), Bandwidth 90% (Hz), as well as the first and the last values of the Peak Frequency Contour (Hz). The selection of these parameters was purely based on their robustness to variation in amplitude and subjectivity in note digitisation across the recordings (Chitnis et al. 2020, Lahiri et al. 2021).

### Statistical Analysis

To analyse the difference in the height of the calling site occupied by different species, we bootstrapped the mean height of the calling site for each species and calculated the 95% CI.

The difference in call frequency between the species was analysed using Principal Component analysis. Principal Component Analysis was performed on the correlation matrix of the eight acoustic parameters in R to reduce the dimensionality.

All the analysis was done using R (Team 2022).

## RESULTS

### Vertical space partitioning between species

There were significant differences in the heights used by the different frog species (Figure 1, *R. akroparallagi* (n=42), *R. ponmudi* (n=31), *P. wynaadensis* (n=24)). While *P. wynaadensis* occupied the lowest height, followed by *R. ponmudi* and then *R. akrolarallagi*. Only two individuals of *R. galndulosus* were found in this study; hence its data was not used in the analysis, but their average height was included in the plot.

**Figure 1:**
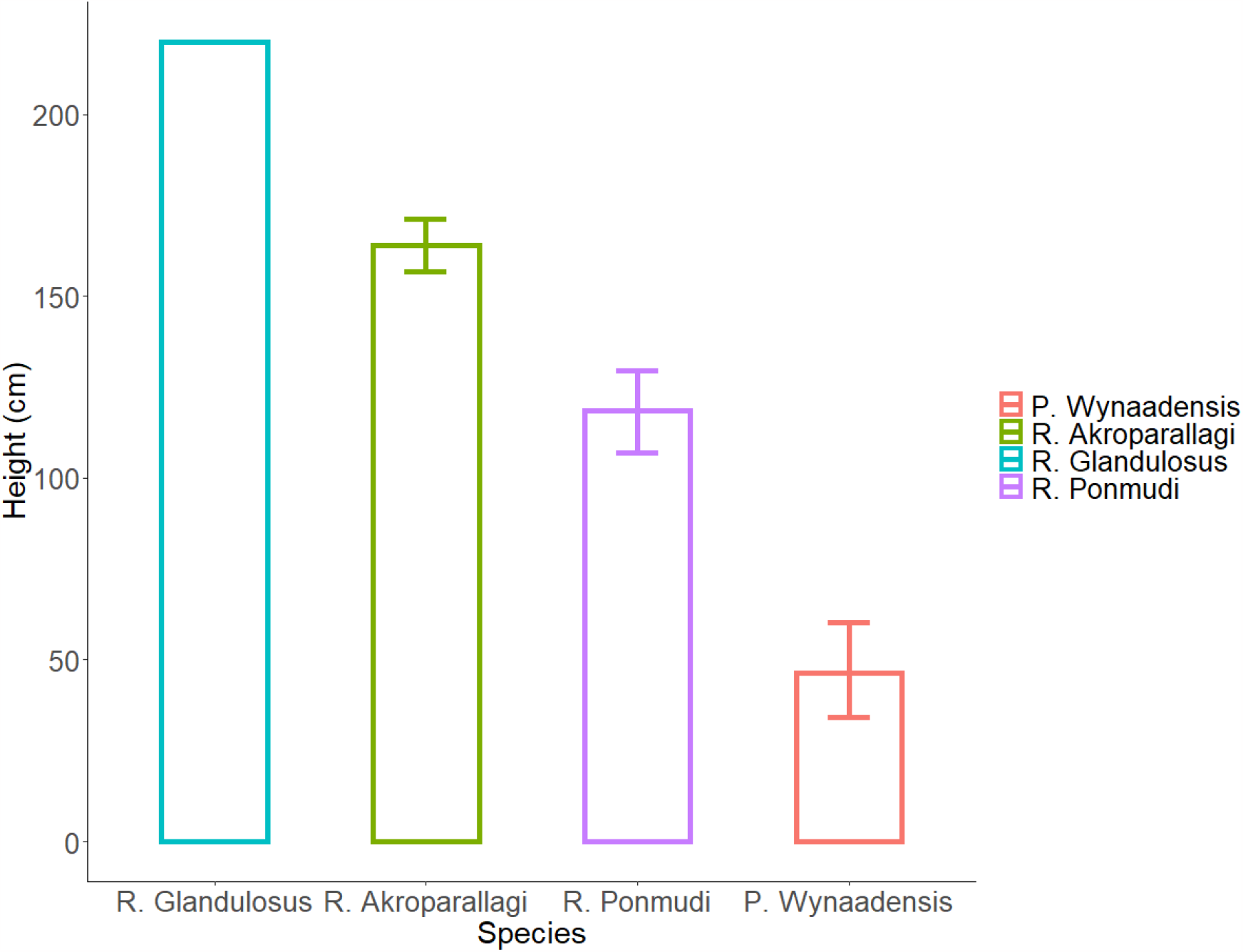
Mean height of the calling sites of the four species of frogs. 95% confidence intervals around the mean generated for three species (*R. galndulosus* (n=2), *R. akroparallagi* (n=42), *R. ponmudi* (n=31), *P. wynaadensis* (n=24)).

### Plant height use, a result of preference

A total of 71 plants with frogs were surveyed, and 74% of the plants hosted only one species, 23.9 % of the plants hosted two species, and only one plant hosted all three species of frogs at the same time. In the single species plants (n=53), each species was found in approximately equal proportions (17, 18, 18 plants were occupied by *R. akroparallagi, R. ponmudi* and *P. wynaadensis*, respectively). This large percentage of single-species occupied plants suggests that each species’ height (mentioned in the previous section of the results) is their preferred height.

### Call frequency segregation among sympatric species

In the Principal Components Analysis (PCA) that summarised the eight acoustic variables (See Methods), the first two Principal Components (PCs) explained 95% of the variation (Supplementary 1). All parameters except for Average Entropy and Bandwidth 90% (Hz) loaded negatively on the PC1.

The principal component space was largely segregated for all species (see Figure 2), suggesting distinct call signatures for the four coexisting species. Interestingly, while we see some overlap between *R. glandulosus* and *P. wynaadensis* (Figure 2), the two species occupy distinctly separate heights (see Figure 1).

**Figure 2:**
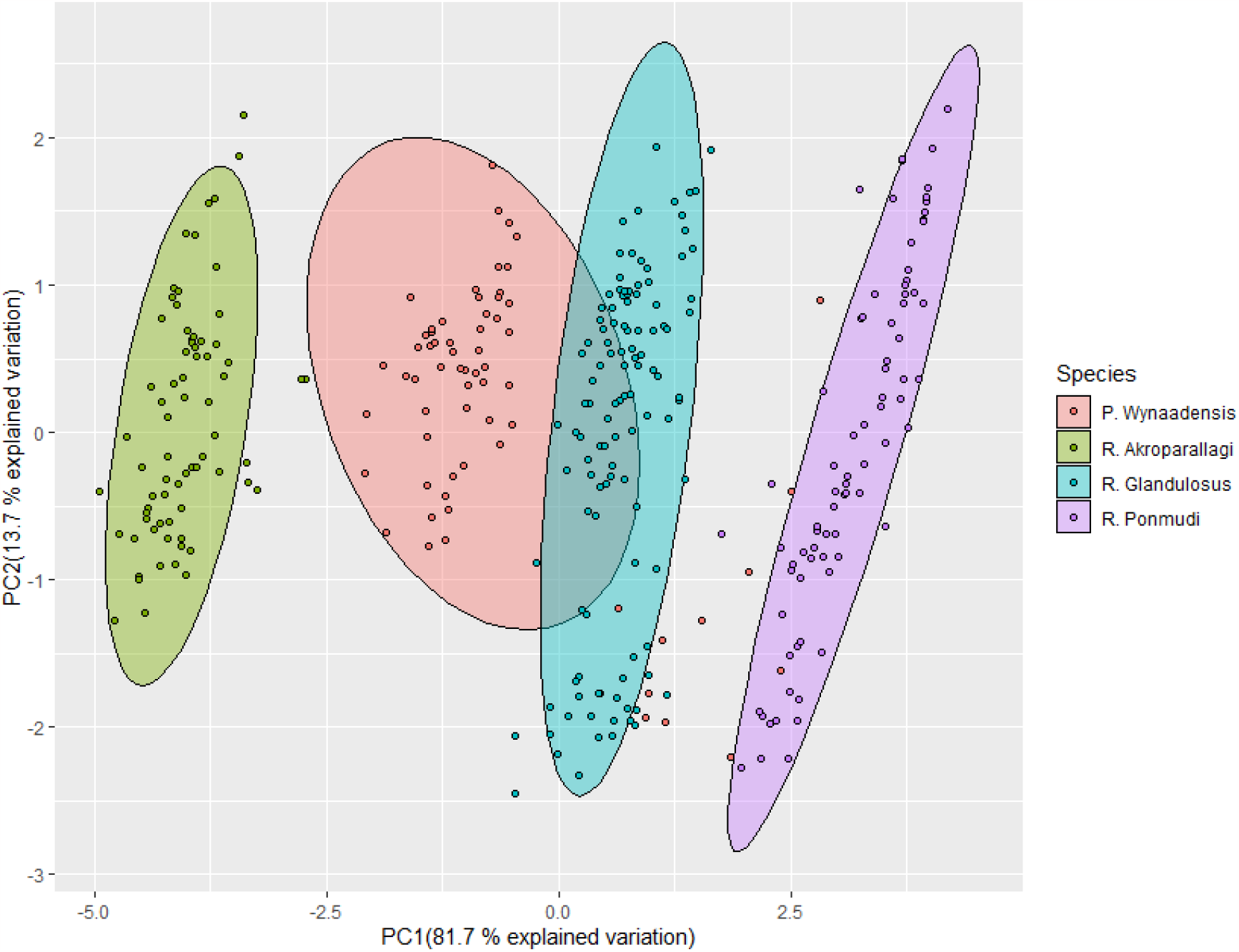
Principal Component Analysis (PCA) plot showing PC1 against PC2 of eight acoustic variables. The lines indicate 95% confidence intervals.

## DISCUSSION

One fundamental question in ecology is understanding how diversity is maintained. For this, ecologists focus on understanding how species coexist in the same area with similar ecology (Chesson 2000). There are various strategies that different species employ to minimise competition and enable coexistence. In vocal species such as frogs, toads, and crickets, differences in the mating call and calling site can play an integral role in conspecific recognition and thus prevent mis-mating. While few empirical studies have tested and established how acoustic and spatial partitioning mechanisms allow coexistence (e.g. Hödl 1977, Lamb 1987, dos Santos Protázio et al. 2015), these mechanisms have not been extensively tested across the globe, particularly in novel habitats.

In endemic tropical amphibian communities such as ours, where coexisting species may be closely related (Vijayakumar *et al*. 2016) and where all species advertise and reproduce in the short breeding season, the consequences of the cocktail party problem may be very strong. Novel habitats such as plantations thus can help us understand how coexisting species solve this problem. By studying both spatial and acoustic differences in a simple and novel habitat, our study attempted to understand the mechanisms of coexistence among four species of bush frogs, which are closely related, breed in the same season, use the same plants for signalling, and vocalise during the same hours.

Our results indicate a clear vertical stratification among the four species of frogs. Interestingly, the frogs spatially segregated with minimal overlap even within the limited vertical space (300cm per coffee plant) (Figure 1). Although each species used or was found occupying a range of heights (Appendix 1), each species clearly had its preference (Figure 1 and Appendix 1). Typically, there is hardly any vegetation between the 300cm high coffee plants and the above canopy tree (tree stands dominated by species such as *Terminalia sp*., *Lagerstroemia microcarpa, Taktona grandis, Dalbergia latifolia, Artocarpus heterophyllus*) due to the annual pruning of the mid-canopy layer to maximise the sunlight availability for the coffee plants (which is linked to coffee productivity). The current spatial distribution and segregation suggest a clear adaptation of these frogs to the management practices in a novel habitat such as coffee plantations.

Vertical stratification has been reported among several arthropods, such as butterflies (Devries *et al*. 1997), fruit flies (Tanabe 2002), ants (Brühl *et al*. 1998), and other herbivore insects (Basset *et al*. 2001). However, the calling site separation among these bush frogs is similar to the calling site preference of other taxa such as cicadas (Sueur 2002) and crickets (Diwakar & Balakrishnan 2007). Although such call stratification is known from some frog communities in the Americas and Europe (Hödl 1977, Lamb 1987, Ptacek 1992, Amézquita *et al*. 2011, Nityananda & Bee 2011, Bignotte-Giró *et al*. 2019, Allen-Ankins & Schwarzkopf 2022), to the best of our knowledge, this is the first evidence from an amphibian community in Asia.

Studies have also investigated horizontal call site partitioning among coexisting species (Schmidt *et al*. 2013). However, our intention of emphasising vertical stratification of space use was not to look at it in isolation but to make sense of it in the context of call masking interference of vocalisations. Horizontal partitioning of call site selection is not expected to greatly reduce masking interference because the signals are still being produced at similar vertical heights and, therefore, would continue to impede information transfer. Also, studying horizontal partitioning is more relevant in a natural habitat where the horizontal space is much larger, unlike in the coffee plantation where a horizontal branch is ∼100cm long.

Our results from call spectral analysis clearly showed the segregation of acoustic trait space among the four species. Even though the calls of these four species of bush frogs are unique (can be clearly distinguished by experienced field researchers), it can be very difficult to differentiate between these species (and between individual males) during a chorus (unless the individual calling is in close vicinity). Similar observations have been made in other studies (e.g. Hödl 1977), and this is the classic cocktail party problem likely faced by both the females (for finding mates) and males (for identifying rivals) of each species in these plantations. So, how do these species overcome this problem?

The solution is revealed by merging the vertical stratification results with the spectral segregation results. In this system, we find that frogs with similar frequency ranges are more spatially separated in space, and species with dissimilar call frequencies occupy adjoining heights on the coffee plants (see Figure 3). This pattern, where we find species with similar frequency ranges tend to segregate spatially, helps avoid acoustic interference and allows the four sympatric species of bush frogs to coexist. Thus, both spatial and spectral segregation are the main mechanisms of coexistence among these frogs during the breeding season. These mechanisms have been suggested to facilitate communicative efficiency in vocal assemblages (Littlejohn 1977, Schmidt *et al*. 2013). The cocktail party problem is indeed a real problem for these species since our preliminary data (unpublished) indicates that all four species vocalise in the early part of the night and in this single (monsoon) season. Future studies could systematically test for temporal partitioning between these species, both within a night (i.e. between dusk and dawn), and along the season.

**Figure 3:**
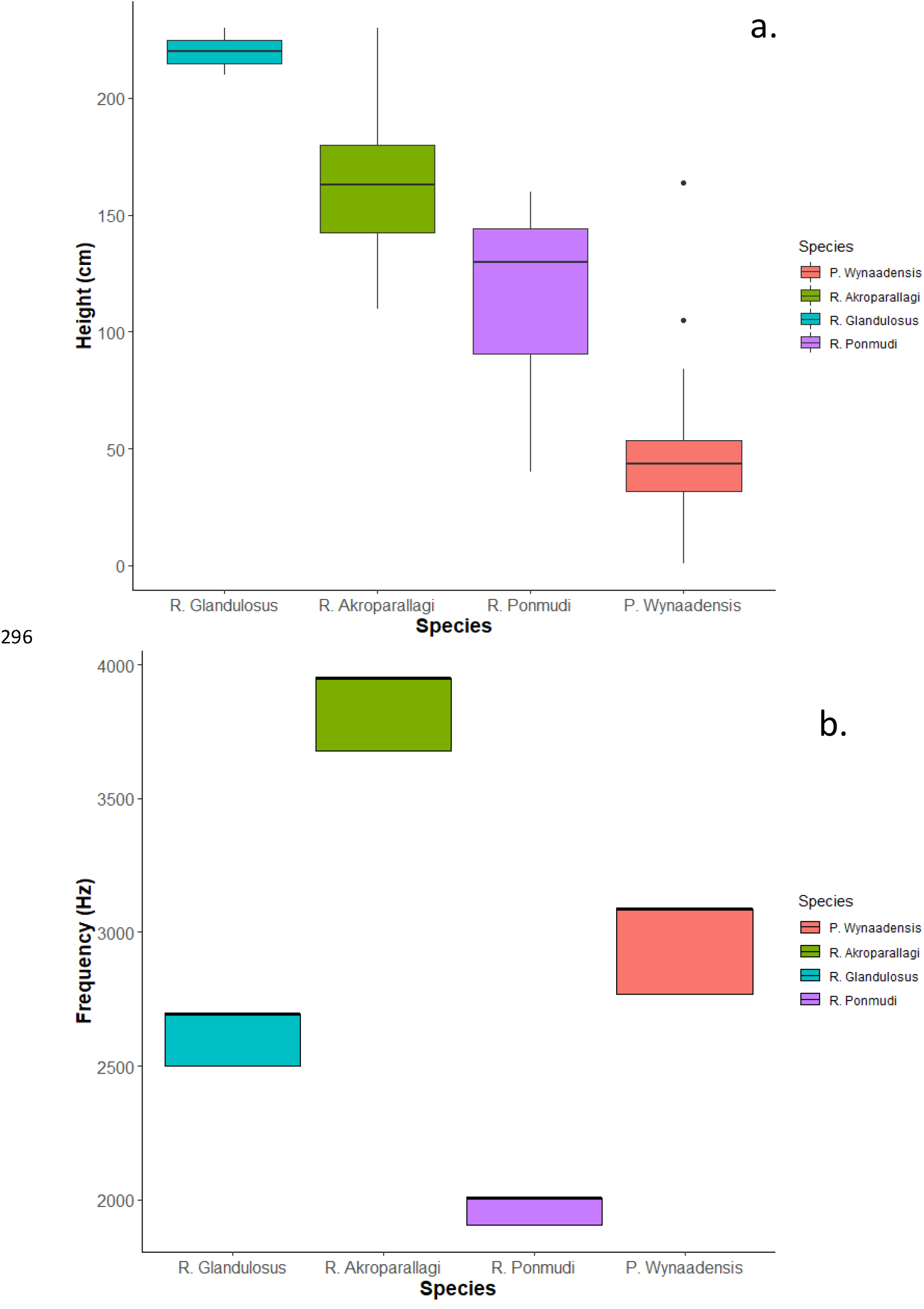
**(a)** Vertical stratification of the calling sites among the four endemic frog species, (**b)** Frequency range occupied by each frog species. The frequency range’s minimum and maximum values denote the mean of the minimum and the maximum Peak Frequency Contours (PFC).

We realise that multiple other factors can also potentially contribute to the frog’s vertical stratification, such as predation risk, choice of background colouration (camouflage), food availability, relative abundances of conspecifics and sympatric species, etc. However, our observed patterns are less likely related to predation risk in this system since snake numbers are very low due to human use. Also, it is less likely that the frog distribution on the plants results from food resource since the frogs are active on the coffee plants only when advertising or seeking mates, usually immediately after a rain shower. Moreover, males of species such as *R. akroparallagi* and *R. glandulosus* produce infrequent calls from the canopy during the intermittent pre-monsoon showers and move down to the coffee plants during monsoon (personal observations). This observed shift in calling site location and spatial segregation during peak breeding season further reinforces that calling site preferences indeed help in reducing masking interference of acoustic signals among these frogs. Future studies could test the importance of other factors, such as background colouration or relative abundance of conspecifics and heterospecifics in microhabitat use. Most importantly, it should be very important to conduct similar studies in natural forested habitats to where they are native. Unfortunately, such studies are missing in this system.

The coffee plantations in Wayanad are highly modified landscapes, with equally spaced and similar-sized (∼ 200-300 cm) coffee plants planted in the shade of tall (>10m high) trees. The coffee plants are maintained at an equal distance from each other, without any overlap, allowing a person to walk between them. Unlike a natural forest, where there is a high amount of heterogeneity, the homogenous nature of plantations, with equal spacing and size of the plants, provided the ideal microhabitat replicates to test these coexistence mechanisms.

## Conclusion

The study showcased how two common and essential coexistence mechanisms allow sympatric male species to minimise competition and enable their coexistence. Data from this field study showed that frogs with similar frequency ranges prefer to stay more spatially separated in space. This helps reduce call interference and allows simultaneous vocalisation of the four species. Apart from this, the interpretation of the obtained results suggests an interesting coexistence strategy in this relatively novel environment. Not only are studies which test both mechanisms in the same system rare, but to the best of our knowledge, this is the first study of its kind on amphibians.

## Acknowledgement

Thanks to Mathan Pilakkavu, Gopalakrishnan P. A, Surendran M, P.C. Chacko, Aboobaker. N and Soman. P for helping find study locations and giving permission to conduct field samplings, and to Azim Premji University for providing funding. AAV would like to thank Manoj Kumar A. V., Sherly Prasannan, and Revathi A. V. for constant support.

## Appendix 1 Additional assessment of vertical space use

To understand the range of the vertical heights being used by the four species, a preliminary study was conducted in the beginning of this study. In this study, visual encounter surveys were conducted by walking 14 transects (35m each) in five farms between 6-11 pm. Each transect was walked four times.

When a male calling frog was encountered, its species was noted, and the height was recorded as a height category (in comparison to the body height of the observer). The six height categories were as follows: 0-10cm (foot–ankle), 10-50cm (ankle-knee), 50-100cm (knee-hip), 100-150cm (hip-shoulder), and 150-200cm (above shoulder), and above 200 cm. This allowed us to understand the frequency distribution of the heights for each bush frog species. A total of 688 individuals [*P. wynaadensis* (n=245), *R. ponmudi* (n=139), *R. akroparallagi* (n=277), *R. glandulosus* (n=21)] were found, and their frequency distribution in the different heights were as follows.

Each frog species occupied a unique height category of the coffee plants (Supplementary Figure 1). While 53% of *P. wynaadnesis* were perched at a height between 10-50 cm (Supplementary Figure 1 a), majority (48%) of the individuals of *R. ponmudi* were perched at height between 100-150 cm (Supplementary Figure 1b). Seventy eight percent of *R. akroparallagi* perched at the 150-200cm height (Supplementary Figure 1c), and 76% of the individuals of *R. glandulosus* perched at height > 200 cm (Supplementary Figure 1d).

**Appendix Figure 1:**
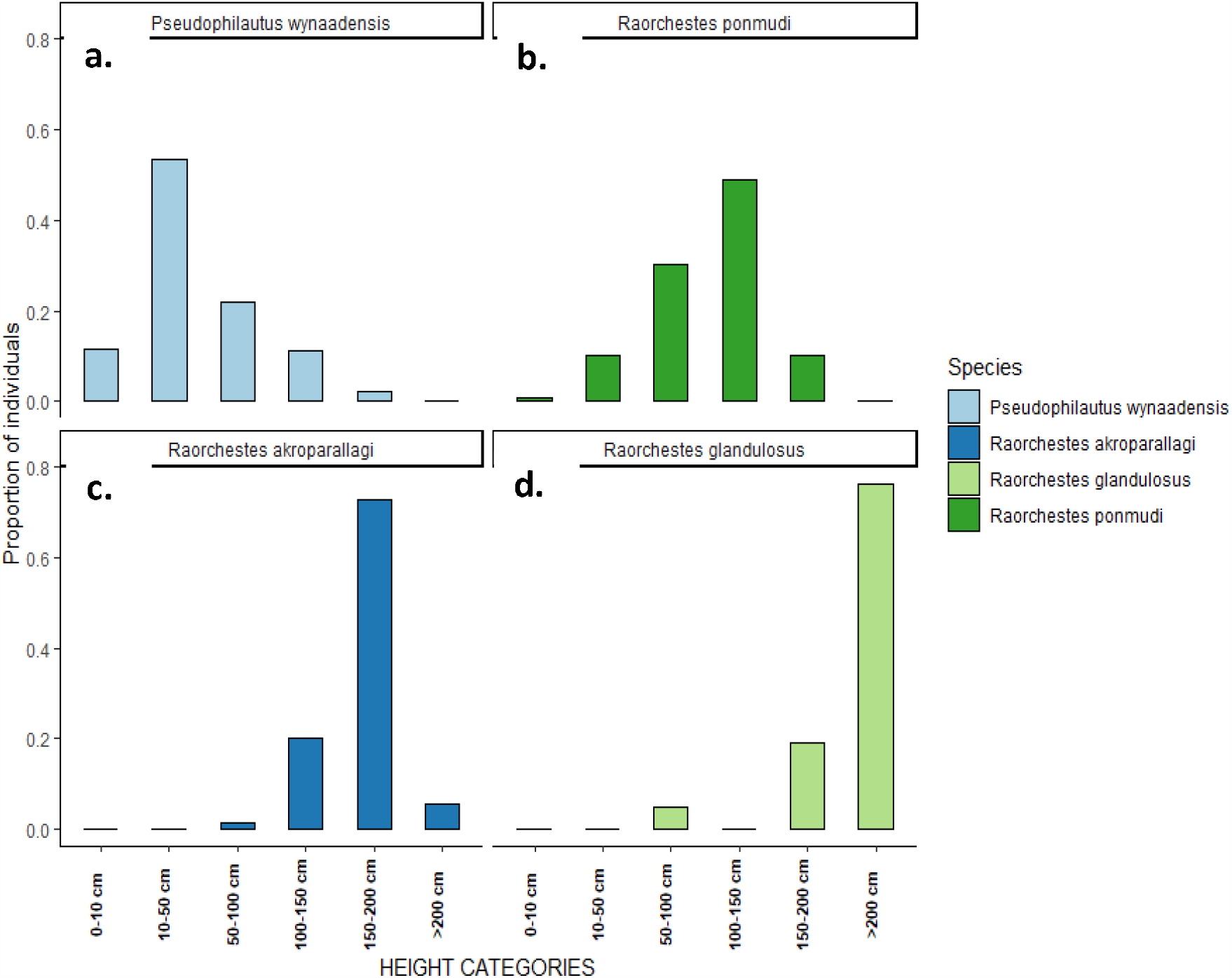
Relative abundance of the four frog species at different height categories from the ground level. (a) *P. wynaadensis* (b) *R. ponmudi* (c) *R. akroprallagi* (d) *R. glandulosus*

**Supplementary Table 1.** Results of the Principal Component analysis for eight acoustic variables. Rows contain factor loadings for each trait, eigenvalues and the proportion of variance explained by each principal component.

|  | PC1 | PC2 | PC3 | PC4 | PC5 | PC6 | PC7 | PC8 |
| --- | --- | --- | --- | --- | --- | --- | --- | --- |
| **Peak Freq (Hz)** | -0.3851 | -0.132 | -0.0510 | 0.2126 | -0.2172 | 0.6408 | -0.0620 | -0.5695 |
| **PFC Max Freq (Hz)** | -0.3873 | 0.0712 | 0.0062 | 0.0642 | 0.7755 | 0.0312 | 0.4878 | -0.0118 |
| **PFC Min Freq (Hz)** | -0.3826 | 0.1560 | -0.1167 | -0.2571 | -0.5350 | -0.3520 | 0.5807 | -0.0456 |
| **Avg Entropy (bits)** | -0.1687 | -0.812 | -0.5579 | -0.0015 | -0.00324 | 0.0098 | -0.0037 | -0.0129 |
| **Center Freq (Hz)** | -0.3859 | 0.1560 | -0.0846 | 0.1170 | -0.1459 | 0.3540 | -0.1108 | 0.8088 |
| **BW 90% (Hz)** | -0.2867 | -0.497 | 0.8115 | -0.0374 | -0.0983 | -0.0085 | -0.0030 | 0.0195 |
| **First PCF value (Hz)** | -0.3851 | 0.1073 | -0.047 | 0.6002 | 0.0061 | -0.5729 | -0.3752 | -0.0924 |
| **Last PCF value (Hz)** | -0.3836 | 0.1293 | -0.0674 | -0.7135 | 0.1843 | -0.1028 | -0.5174 | -0.100 |
| **Eigenvalue** | 2.5577 | 1.0472 | 0.5374 | 0.1717 | 0.1354 | 0.1214 | 0.0827 | 0.0564 |
| **% Explained variance** | 81.77 | 13.71 | 3.61 | 0.361 | 0.229 | 0.184 | 0.086 | 0.040 |
| **Cumulative % explained** | 81.77 | 95.48 | 99.09 | 99.46 | 99.69 | 99.87 | 99.96 | 100 |

